# Asymmetric Diversification of Mating Pheromones in Fission Yeast

**DOI:** 10.1101/366260

**Authors:** Taisuke Seike, Chikashi Shimoda, Hironori Niki

**Affiliations:** Genetics Strains Research Center, National Institute of Genetics, 1111 Yata, Mishima, Shizuoka, 411-0806, Japan; Graduate School of Science, Osaka City University, 3-3-138 Sugimoto, Sumiyoshi-ku, Osaka, 558-8585, Japan

## Abstract

In fungi, mating between partners critically depends on the molecular recognition of two peptidyl mating pheromones by their respective receptors. The fission yeast *Schizosaccharomyces pombe* has two mating types, Plus (P) and Minus (M), which secrete two different mating pheromones: P-factor recognized by Mam2, and M-factor recognized by Map3, respectively. Our recent study demonstrated that a few mutations in both M-factor and Map3 can trigger reproductive isolation, a cause of speciation, in *S. pombe*. Here we explored the mechanism underlying reproductive isolation through genetic changes of pheromones and receptors. We investigated the diversity of genes encoding the pheromones and their receptor in 150 *S. pombe* wild strains. Whereas the amino acid sequences of M-factor and Map3 were completely conserved, those of P-factor and Mam2 were very diverse. In addition, the P-factor gene contained varying numbers of tandem repeats of P-factor (4–8 repeats). We also explored the recognition specificity of pheromones between *S. pombe* (Sp) and its close relative *Schizosaccharomyces octosporus* (So). So-M-factor did not have an effect on *S. pombe* P-cells, but So-P-factor had a partial effect on *S. pombe* M-cells, allowing them to mate successfully. Thus, recognition of M-factor seems to be tight, whereas that of P-factor is relatively loose. Moreover, diversity of P-factor and Mam2 might be due to a P-factor-specific peptidase. Overall, the asymmetric system for pheromone recognition in yeasts seems to allow flexible adaptation to mutational changes in the combination of pheromone and receptor while maintaining tight recognition for mating partners.

## Introduction

New species may emerge when two populations can no longer interbreed. Since the time of Charles Darwin, the processes by which new species originate from previously interbreeding individuals have remained highly controversial^1^. Reproductive isolation, which restricts gene flow between sympatric populations, is one of the key mechanisms of speciation^2^. Species become reproductively isolated by mainly one of two barriers. The first barrier, termed prezygotic isolation, prevents interbreeding of closely related species. Prezygotic isolation is mainly caused by changes in signals that enable individuals to appropriately recognize the opposite sex: for example, pheromones in insects^3,4^ and amphibians^5,6^; body color in fish^7^; and song in birds^8^. The second barrier, termed postzygotic isolation, is hybrid sterility or nonviability of the descendants of related species^9^. In postzygotic isolation, the hybrids are incompatible owing, for example, to aneuploidy and chromosomal rearrangement in plants^10,11^. Although both types of reproductive isolation have been frequently studied in higher organisms, far less is known about them in fungi^12^.

In ascomycetes fungi including yeasts, mating between partners critically depends on the molecular recognition of peptidyl mating pheromones by receptors^13-16^. Our recent study in the fission yeast *Schizosaccharomyces pombe* demonstrated that several mutations of a pheromone and its corresponding receptor create a prezygotic barrier that can give rise to a new species^15^. This experimental observation supports the idea that, first, pheromone–receptor systems drive reproductive isolation through very subtle variations in nature; and second, a new prezygotic barrier might potentially lead to the origin of a new species. Thus, genetic alterations of pheromones and their receptors are likely to be important to promote speciation in yeasts. More generally, however, loss of pheromone activity may result in extinction of an organism’s lineage; therefore, coevolution of pheromones and their receptors might occur gradually and/or coincidently before speciation happens. This hypothesis is an attractive explanation for prezygotic isolation in yeasts; however, the mechanisms of ongoing reproductive isolation through genetic alterations of pheromone–receptor systems in nature remain to be elucidated.

*S. pombe* has two mating types, Plus (P) and Minus (M)^17-19^. Under nitrogen-limited conditions, two haploid cells of opposite mating types mate via reciprocal stimulation of their mating pheromone receptors^20^. On the one hand, P-cells secrete a mating pheromone called P-factor, a simple 23-amino acid peptide, which is recognized by its corresponding G-protein coupled receptor (GPCR) Mam2 on M-cells^21,22^. The *map2*^+^ gene encodes a precursor polypeptide containing four tandem repeats of mature P-factor. P-factor is secreted by the standard secretory pathway^22^. On the other hand, M-cells secrete a mating pheromone called M-factor, which is recognized by its GPCR Map3 on P-cells^23^. Mature M-factor, a farnesylated and methylated peptide of nine-amino acids, is encoded by three redundant genes: *mfm1*^+^, *mfm2*^+^, and *mfm3*^+24,25^. Polypeptides produced from each of these *mfm* genes ultimately yield one copy of M-factor. M-factor is secreted specifically by the ATP-binding cassette (ABC) transporter Mam1^26-28^. *S. pombe* M-cells also produce a pheromone-degrading enzyme encoded by the *sxa2*^+^ gene^29,30^. Sxa2 is a serine carboxypeptidase that specifically degrades extracellular P-factor. The C-terminal residue (Leu) of P-factor is removed by Sxa2 outside the cells^30^, and the resulting P-factor lacking Leu is inactive and not recognized by Mam2^31^. Yeast cells sense a gradient of pheromones secreted by the opposite cell and then extend a mating projection towards the pheromone source^32^. Degradation of the pheromones by the peptidase is thought to make the gradient more stable. By contrast, an enzyme that degrades M-factor has not yet been found. Instead, expression of the *mfm* genes encoding M-factor might be differentially controlled to facilitate fine-tuning. These differences in the two mating pheromones, including chemical structure, secretion pathway, and degradation, are widely common in ascomycetes^13^, but the biological significance remains unclear.

We considered that the redundancy in the pheromones might allow unrestricted diversification. Indeed, the standard laboratory strain of *S. pombe*, L968, has four copies of P-factor in the corresponding gene and three genes encoding M-factor^33^. Thus, evolution has a mechanism for creating new versions of pheromones, while at the same time retaining the ability to mate via the original versions. In this study, therefore, we examined how pheromones and their receptors coevolve in nature. First, we examined pheromone diversity by determining the nucleotide sequences of the pheromone and receptor genes in 150 non-standard *S. pombe* strains^34, 35^, finding that the amino acid sequence of M-factor was completely conserved, whereas that of P-factor was very diverse. In some strains, for example, the copy number of P-factor increased to 4–8 repeats. Second, we analyzed the specificity of pheromones–receptor recognition between *S. pombe* (Sp) and the related species *Schizosaccharomyces octosporus* (So). Whereas So-M-factor was not functional in *S. pombe*, all So-P-factors tested were partially functional in *S. pombe*, enabling these cells to mate successfully using So-P-factors. Thus, recognition of M-factor is highly specific, whereas that of P-factor is relatively loose. Probably, the asymmetric system for pheromone recognition in yeast allows flexible adaptation to mutational changes in a combination of pheromones and receptors, while maintaining tight recognition of mating partners.

## Results

### The amino acid sequences of M-factor and Map3 are completely conserved, whereas those of P-factor and Mam2 are diverse

To investigate the diversity of *S. pombe* mating pheromones and their corresponding receptors in nature, we collected 150 wild strains whose origins differ from that of the standard laboratory strain, L968 (Supplementary Tables 1 and 2), first described by U. Leupold^33^. These strains were derived from various countries and regions (Fig. 1a), and were isolated from several different sources (Supplementary Table 2)^34^. We sequenced the M-factor genes (*mfm1, mfm2*, and *mfm3*), the receptor gene for M-factor (*map3*), the P-factor gene (*map2*), and the receptor gene for P-factor (*mam2*) of all 150 strains, and compared the nucleotide sequences with that of the L968 strain registered in the database (PomBase; https://www.pombase.org). We constructed a phylogenetic tree based on the sequences of these six genes in the 151 *S. pombe* strains (Fig. 1b). Many nucleotide differences in these genes were found among the strains (Supplementary Tables 3 and 4). As shown in Fig. 1b, the sequence patterns of these strains were relatively diversified, with characteristic patterns depending, in part, on region (*i.e*., Europe or America).

**Table 1.** Summary of polymorphisms of the seven pheromone-associated genes in 151 *S. pombe* strains.

| Gene |  | Polymorphism | Nucleotide sequence pattern<sup>a</sup> |
| --- | --- | --- | --- |
| *mfm1*<br>(M-factor) | 1 | WT | WT, A1, A2, A3, A4, A5, A6, A7 |
| *mfm2*<br>(M-factor) | 1 | WT | WT, B1, B2, B3, B4, B5 |
| *mfm3*<br>(M-factor) | 1 | WT | WT, C1, C2, C3 |
| *map2*<br>(P-factor) | 6 | WT (P2) | WT, D1, D2, D3, D4, D5, D6, D7, D8, D9, D10, D11, D12, D13, D14, D15 |
|  |  | T1S, A8V, V16A (P1) | WT, D1, D2, D3, D4, D5, D6, D7, D8, D9, D10, D12, D13, D14, D15 |
|  |  | T1S, Q10H (P3) | WT, D2, D3, D4, D5, D6, D7, D8, D9, D10, D11, D12, D13, D14, D15 |
|  |  | T1S, A8V (P4) | D1, D3, D5, D6, D8, D10, D11 |
|  |  | P18L (P5) | D9 |
|  |  | Q10H (P6) | D13 |
| *map3*<br>(M-factor receptor) | 1 | WT | WT, E1, E2, E3, E4 |
| *mam2*<br>(P-factor receptor) | 5 | WT | WT, F4 |
|  |  | I101V | F2 |
|  |  | H155R | F1 |
|  |  | K158N | F5 |
|  |  | T164S | F3 |
| *sxa2*<br>(P-factor-degrading enzyme) | 4 | WT | WT |
|  |  | K62E, N82D, Q184E, N208Y, V248M | G1, G2, G3, G5 |
|  |  | K62E, N82D, Q184E, N208Y, V248M, G315D | G4, G6 |
|  |  | K62E, N82D, Q184E, N208Y, V248M, V321I | G7 |

<sup>a</sup>These patterns are described in Supplementary Table 3.

The extent of degradation of each P-factor was determined by the amount of leucine released. Synthetic P-factor peptide (200 μM) was mixed with cell-free culture supernatant (total protein 100 ng), either containing active Sxa2 (TS407 (Sxa2^+^)) or lacking Sxa2 (TS406 (Sxa2^-^)) in 50 mM citrate buffer, pH 5.5. All reactions were performed at 30°C for the indicated times (10 and 60 minutes) with gentle shaking, and were stopped by 0.5% trifluoroacetic acid. Leucine was measured by a branched-chain amino acids (leucine) kit. Data that were calculated by taking a difference between two values in the presence/absence of Sxa2 are the mean ± S.E.M. of at least triplicate samples. ‘P2-Leu’ is a 22-amino acid peptide whose C-terminal Leu residue of P2 was removed.

**Figure 1.**
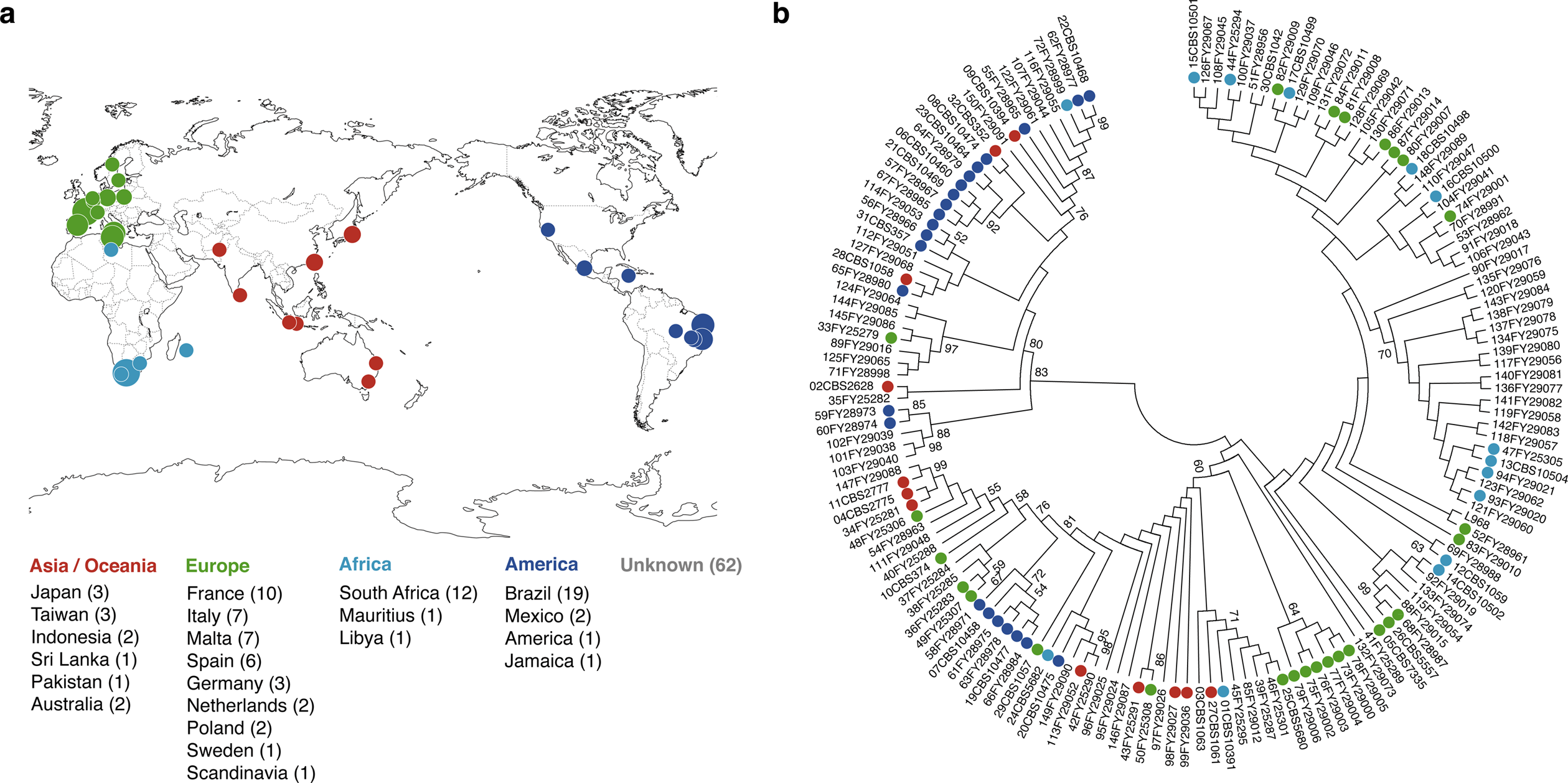
An overview of *S. pombe* wild strains used in this study. **a**, Geographical origins of the 150 wild strains (Supplementary Table 2). Colored circles indicate the approximate countries and regions of the isolated strains, which are color-coded by continents: Asia/Oceania, Europe, Africa, and America. Circle size increases in proportion to the number of strains isolated from each region; the actual number of strains is shown in parentheses. Location was unknown for 62 strains. **b**, Phylogenetic tree of 151 *S. pombe* strains including the laboratory strain (L968). The analysis involved combined multiple sequence alignments for *mfm1, mfm2, mfm3, map2, map3*, and *mam2*. Evolutionary history was inferred by using the Neighbor-Joining^59^. The optimal tree with the sum of branch length = 0.03655487 is shown. The evolutionary distances were computed by using the Kimura 2-parameter method^60^, and are shown in units of the number of base substitutions per site. All ambiguous positions were removed for each sequence pair. There were a total of 3,737 positions in the final dataset. Evolutionary analyses were conducted in MEGA7^61^.

Notably, all three *mfm* genes of all 151 strains (i.e., 453 genes in total) produced an identical mature M-factor peptide, YTPKVPYMC^Far^-OCH3, (102 genes have been previously reported^36^). Many mutations were found in the pro-sequences and introns, but one exceptional mutation in the mature M-factor-encoding region was a synonymous change, causing no amino acid substitution (Supplementary Table 4). Moreover, the amino acid sequence of Map3 in the 150 strains was also identical to that in the L968 strain (Table 1). In other words, the amino acid sequences of the M-factor/Map3 pair seem to be completely conserved in nature. By contrast, the sequences of the *map2* genes were very diverse (Supplementary Tables 3 and 4). Whereas the *map2* gene in L968 carries four P-factor-encoding tandem repeats (P1-P2-P3-P2), extensive variations in the number of repeats were observed in the 150 strains, ranging from 4 to 8 (Fig. 2a). In addition, the *map2* genes of the wild strains were predicted to produce six different mature P-factor peptides (P1–P6; see Fig. 2a). Interestingly, there were five different amino acid sequences of Mam2 across the 150 strains (Table 1). Thus, the amino acid sequences of the P-factor/Mam2 pair seem to be very diverse in nature. Collectively, these findings show that the two mating pheromones in *S. pombe* have diversified asymmetrically.

**Figure 2.**
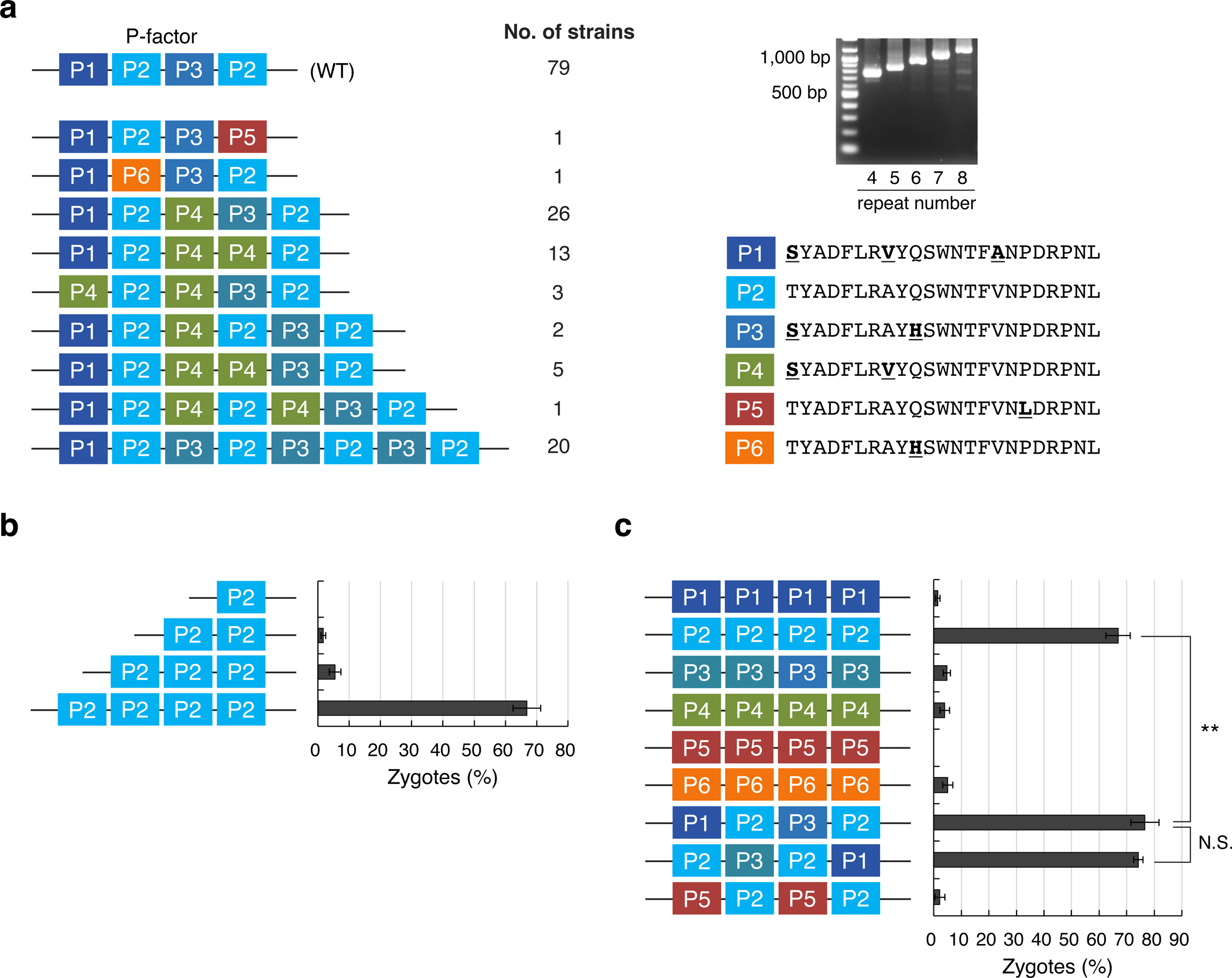
Six different P-factors are found in nature. **a**, Diversified pattern of the *map2* in nature. In the laboratory strain (WT; L968), the *map2*^+^ gene carries four tandem P-factor-encoding repeats (P1-P2-P3-P2). About half of the 150 wild strains had a *map2* gene with a higher number of P-factor repeats (5-8 repeats). The electrophoresis gel shows PCR products amplified with the primer set oTS85/86 from the genomic DNA of representative strains. The six different P-factors (P1–P6) found in nature are shown below. P2 peptide is the standard for P-factor of *S. pombe;* the amino acids that differ from the P2 peptide are underlined in bold. **b**, Zygote frequency of strains with a *map2* ORF containing different numbers of P-factor-encoding repeats (*i.e*., 1-4 identical repeats of the P2 peptide). **c**, Zygote frequency of strains producing various P-factors (all with 4 repeats). At least 300 cells were examined for each sample. Data are the mean ± S.E.M. of triplicate samples. Statistical significance was assessed by *t*-test (\*\**p* <0.01; N.S., not significant).

### Wild *S. pombe* strains simultaneously produce multiple P-factor peptides

Having observed increased numbers of P-factor-encoding repeats in the *map2* gene in about half of the 150 strains (Fig. 2a), we tested whether repeat number directly affects mating frequency. The native 4-repeat Map2 open reading frame (ORF) from the L968 strain was replaced with an ORF carrying different numbers of the P2 repeat (see Methods). We found that a decrease in P2 repeat number (<4 repeats) resulted in an extremely low frequency of zygotes (Fig. 2b). For example, the strain with the 3-repeat ORF produced less than one-tenth of the zygotes (%) of the strain with the 4-repeat ORF (Fig. 2b). However, few significant differences in mating efficiency were observed among the strains carrying an ORF with more than 4 repeats (Supplementary Fig. 1a).

To compare the activity of the different P-factor peptides (P1–P6), we introduced a modified *map2* gene carrying four tandem repeats of each P-factor into the P-factor-less strain (FY23418; Supplementary Table 1), in which the native *map2* gene had been deleted (see Methods). The resulting strains each produced one of the six P-factors. The mating efficiency of these strains was assessed by the frequency of zygotes. The strain producing four P2 peptides from the *map2* gene showed a high frequency of zygotes (66.9 ± 4.4%); surprisingly, however, the zygote frequency of the remaining strains producing the other P-factor peptides (P1, P3–P6) was extremely low or zero (Fig. 2c). Furthermore, in a strain in which the native *map2* gene carrying a sequence of repeats (P1-P2-P3-P2) was introduced, the frequency of zygote was fairly high (76.6 ± 5.1%), despite the production of only two copies of the P2 peptide (Fig. 2c). To determine whether the order of nucleotide sequences encoding mature P-factor affects zygote frequency, we also introduced a *map2* gene carrying a permuted sequence of repeats (P2-P3-P2-P1) into the FY23418 strain. Similar to the strain with the native sequence, this strain also mated at high frequency (74.2 ± 1.7%) (Fig. 2c); therefore, the effect of peptide-coding position was negligible.

Next, we considered that, if mature P-factor production is modulated by a length-dependent biosynthetic pathway, the low mating efficiency observed in strains carrying an ORF with fewer than 4 repeats might be attributed to translation level. To examine this possibility, we constructed a *map2* gene carrying the four tandem P-factor-encoding repeats P5-P2-P5-P2, and introduced it into the FY23418 strain. Surprisingly, however, this strain was almost sterile as the strain with the 2-repeat ORF (Fig. 2c). Thus, the apparently inactive P1 and P3 peptides might have important roles during the mating process. The amino acid sequences of the P1 and P3 peptides slightly differ from that of the P2 peptide (Fig. 2a). In fact, some peptides with slight mutations of P1–P3 had markedly decreased mating efficiency (Supplementary Fig. 1b). These results indicate that the substitution of only a few amino acids of P-factor might have a large influence on copulation, and that the P2 peptide is likely to be most compatible with the native Mam2 receptor; curiously, however, all of the wild *S. pombe* strains simultaneously produce multiple P-factor peptides (such as P1 and P3).

### Production of multiple P-factors might be advantageous for mate choice

As described above, the *map2* gene was found to contain at least four tandem repeats encoding multiple P-factor peptides in all 150 wild strains. To assess the effect of producing multiple P-factors, we performed a quantitative competitive mating assay. In this experiment, P-cells producing a multiple P-factor (P1-P2-P3-P2) and P-cells producing a single P-factor (P2-P2-P2-P2) were inoculated with wild-type M-cells onto MEA at a cell number ratio of 1:1:2. The three strains were differentially marked by different drug-resistant markers (see Methods). After incubation for 24 hours, the cell suspension was spread onto YEA plates containing combinations of the appropriate drugs. Doubly resistant hybrid descendants of mating between P-cells and M-cells were counted to determine the recombinant frequency.

The assay indicated that M-cells showed a slight preference to mate with P-cells producing a multiple-type P-factor as compared with a single-type P-factor (Fig. 3). This tendency was not influenced by the type of drug-resistant marker used (Fig. 3). These results suggest that the production of multiple P-factors might be advantageous for mate choice by M-cells, even though the relative proportion of high-active P2 peptide is lower.

**Figure 3.**
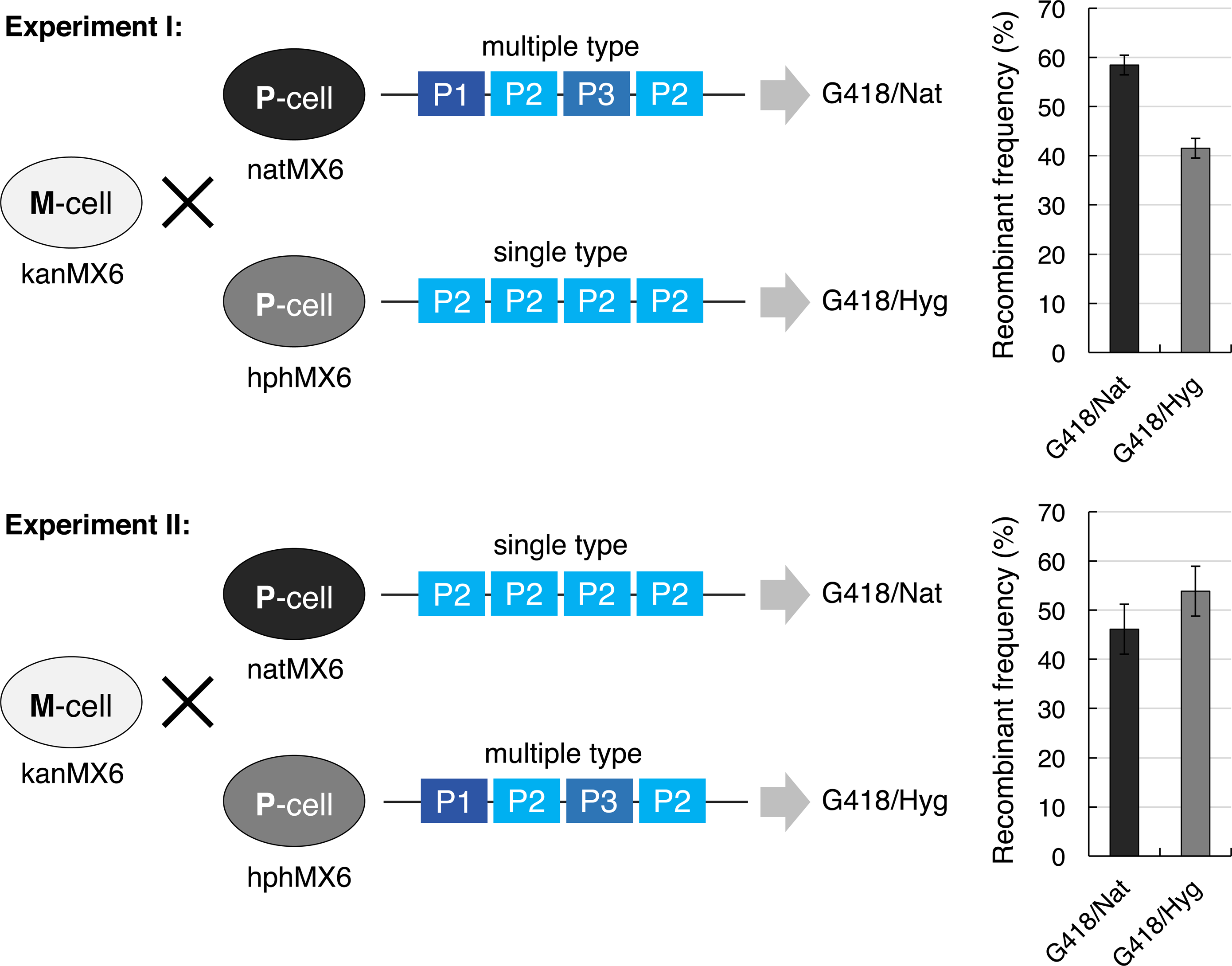
Recombinant frequency of strains producing a multiple or single type of P-factor. The assay was conducted by using heterothallic haploid strains carrying a kanMX6, hphMX6, or natMX6 drug-resistant marker as follows: Experiment I, wild-type M-cells (FY23362; kanMX6), P-cells producing a multiple-type P-factor (FY23424; natMX6), and P-cells producing a single-type P-factor (TS232; hphMX6); Experiment II, wild-type M-cells (FY23362; kanMX6), P-cells producing a single-type P-factor (TS59; natMX6), and P-cells producing a multiple-type P-factor (TS200; hphMX6). The same number of two P-cells and twice the number of cells of the M-cells were mixed and grown on MEA. After incubation for 1 day, the cultures were diluted, and then spread onto YEA containing 100 μg/ml of the following drugs: YEA+G418/Nat and YEA+G418/Hyg. After 3 days of incubation, the YEA plates were photographed. The colony numbers on plates were counted and the recombinant frequency was calculated. Data are the mean ± S.E.M. of three independent experiments.

### Most P-factor peptides are recognized by Mam2

In *S. pombe*, P-factor is largely degraded outside the cell by the carboxyl peptidase Sxa2^30^. Next, therefore, we examined whether each P-factor peptide is truly recognized by Mam2 in the absence of Sxa2. An M-type strain lacking Sxa2 (TS402; Supplementary Table 1) was treated with synthetic P-factors at different concentrations (0–1000 nM) in nitrogen-free liquid medium (EMM2-N). Yeast cells elongate a mating projection (‘shmoo’) when they sense sufficient pheromones; therefore, the ability of Mam2 to recognize the different P-factors was determined by measuring the ratio between the length (L) and width (W) of an individual cell (see Methods). We defined cells with a L/W ratio of 2.0 or more as “shmooing” cells.

The addition of P2 peptide clearly induced cell elongation at 10 nM after 1 day of incubation. According to our definition, one third of cells treated with P2 peptide (10 nM) became shmooing cells in response to P-factor after 1 day (Fig. 4). Notably, cells treated with 10 nM P1, P3, P4, and P6 also elongated, indicating that these peptides were recognized by Mam2. As compared with P2 peptide, however, the proportion of shmooing cells after treatment with these peptides was relatively low (15%–23% at 10 nM; see Fig. 4). By contrast, the P5 peptide, produced only by the 24CBS5682 strain (Fig. 2a and Supplementary Tables 3 and 4), was not recognized by Mam2 even when the cells were treated with a concentration of 1000 nM (Fig. 4). In conclusion, the shmoo formation assay indicated that most P-factor peptides are sufficiently recognized by Mam2 *in vitro*.

**Figure 4.**
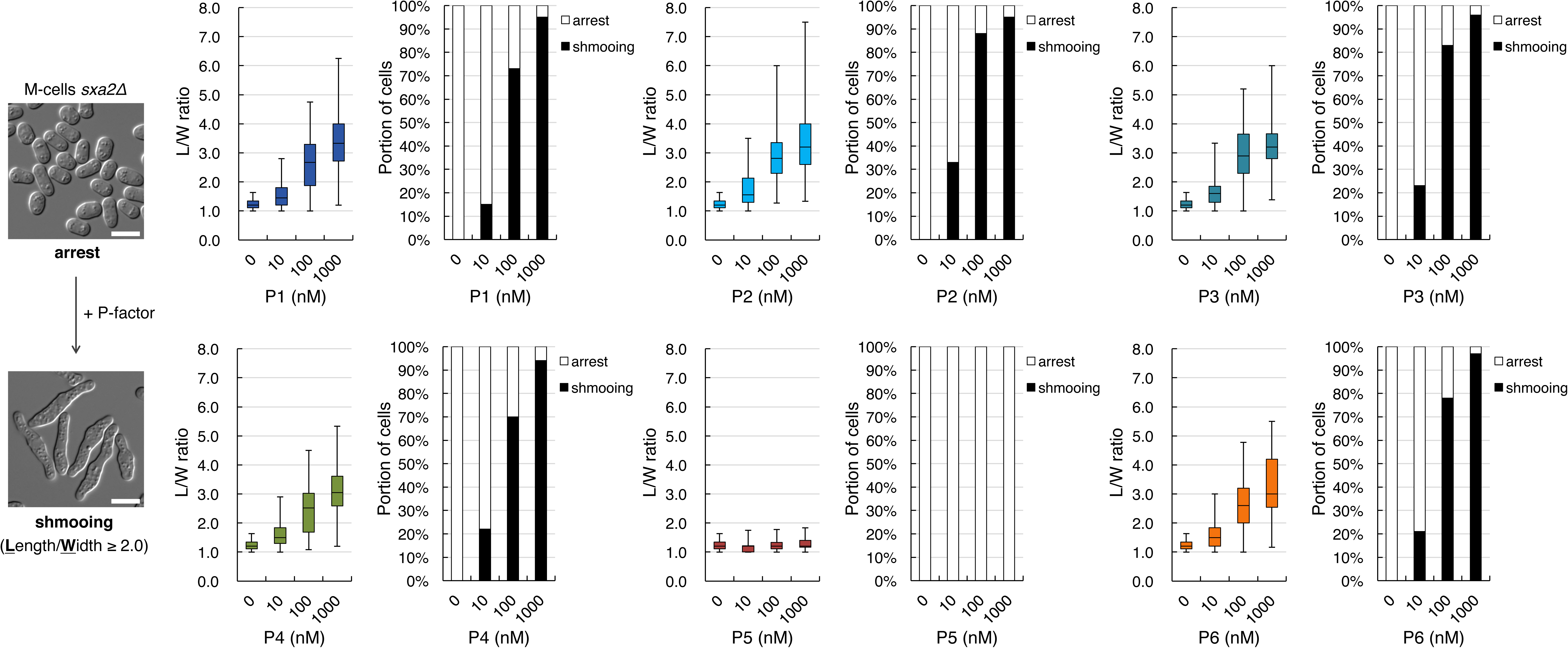
Shmooing assay of synthetic P-factor peptides. M-cells lacking the *sxa2*^+^ gene (TS402) treated with synthetic P-factor at different concentrations (0, 10, 100, and 1000 nM) were incubated in EMM2-N medium with gentle shaking for 24 hours. The ability of each P-factor peptide to induce shmooing was assessed by the length/width (L/W) ratio of a cell. Cells with an L/W ratio of 2.0 or more were defined as shmooing cells (shown in black); those with a ratio of 2.0 less were defined as arrested cells (shown in white). Box-and-whisker plots reperesent the distribution of the L/W ratio; for each peptide, at least 100 cells each were measured. Scale bar, 5 μm.

### P2 peptide is degraded more rapidly by Sxa2

Sxa2 removes the C-terminal Leu residue of P-factor. Based on the above results, we considered that differences in the processing of each P-factor peptide by Sxa2 might affect pheromone-based mate choice. To examine this possibility, the *sxa2*^+^ ORF was cloned into the pTS111 plasmid downstream of the *nmt1*^+^ promoter, which is strongly expressed in absence of thiamine^37^ (see Methods). The resulting plasmid (pTS284) was then introduced into the TS402 strain lacking Sxa2. The resulting cells were grown in EMM2 medium without thiamine to induce the *nmt1*^+^ promoter, and the culture supernatant was assayed to confirm carboxypeptidase activity (see Methods). Next, each P-factor peptide (200 μM) was mixed with an aliquot of the cell-free culture supernatant including abundant active Sxa2 (total protein: 100 ng) for up to 60 minutes, and the leucine concentration was determined as a measure of Sxa2 activity.

All six P-factor peptides (P1–P6) were completely degraded after 60 minutes in the presence of Sxa2 (Table 2). Unexpectedly, degradation of P2 peptide was found to be more efficient than that of other P-factor peptides. In 10 minutes, approximately 70% of P2 peptide (133.0 ± 21.7 μM) was degraded, as compared with, for example, ~50% of P1 (97.7 ± 5.8 μM; see Table 2). These results suggest that native Sxa2 might not efficiently degrade P-factor peptides containing a few residues that differ from P2 peptide. Thus, it is possible that cells produce multiple P-factor peptides in order to escape degradation by Sxa2, and thereby improve their likelihood of selection by M-cells during mating.

**Table 2.**
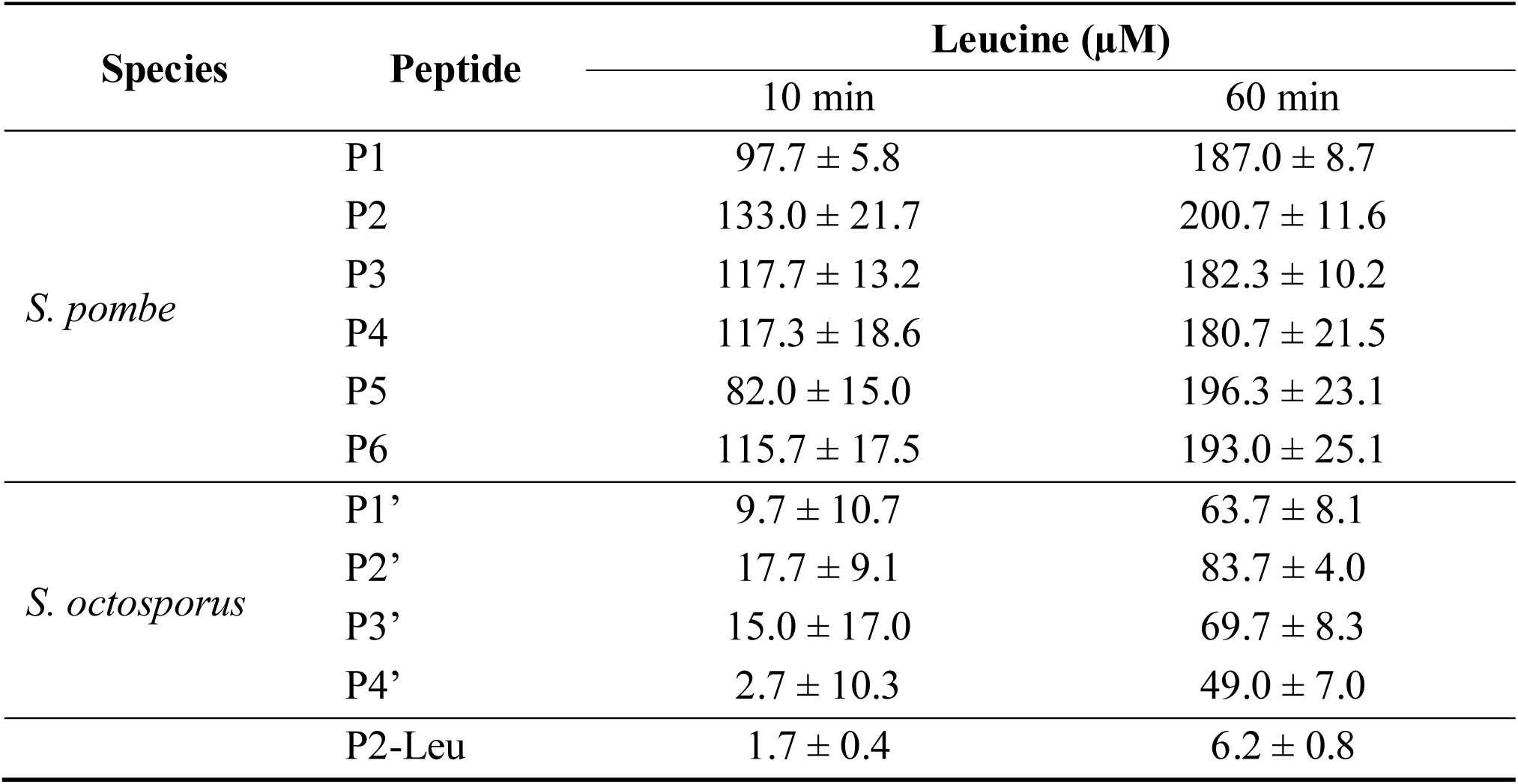
Amount of leucine released from P-factor by Sxa2 of *S. pombe*.

### Asymmetric diversity of two pheromones is seen in closely related species

As described above, the amino acid sequences of M-factor were completely conserved, whereas those of P-factor were diversified in 150 *S. pombe* wild strains analyzed. To determine whether this asymmetry in pheromone diversity is common to other species, we analyzed the nucleotide sequences of both M-factor and P-factor genes in *Schizosaccharomyces octosporus*, the species most closely related to *S. pombe*.

Whole-genome sequences of *S. octosporus* have been determined by the Broad Institute^38^, and indicate that this species has six M-factor-encoding genes (hereafter called ‘*So-mfm1-So-mfm6*’) and a P-factor-encoding gene (hereafter called ‘ *So-map?*). The six redundant genes (*So-mfm1-So-mfm6*) encode M-factor peptides with the same amino acid sequence (Fig. 5a); therefore, the primary structures of the putative So-M-factors are the same, YQPKPPAMC^Far^-OCH_3_ (Fig. 5a). By contrast, the *So-map2* gene carries seven tandem So-P-factor repeats, which encode four different So-P-factors (Figs. 5b and 6c), similar to *S. pombe*. Interestingly, therefore, the two pheromones have also diversified asymmetrically in this closely related species.

**Figure 5.**
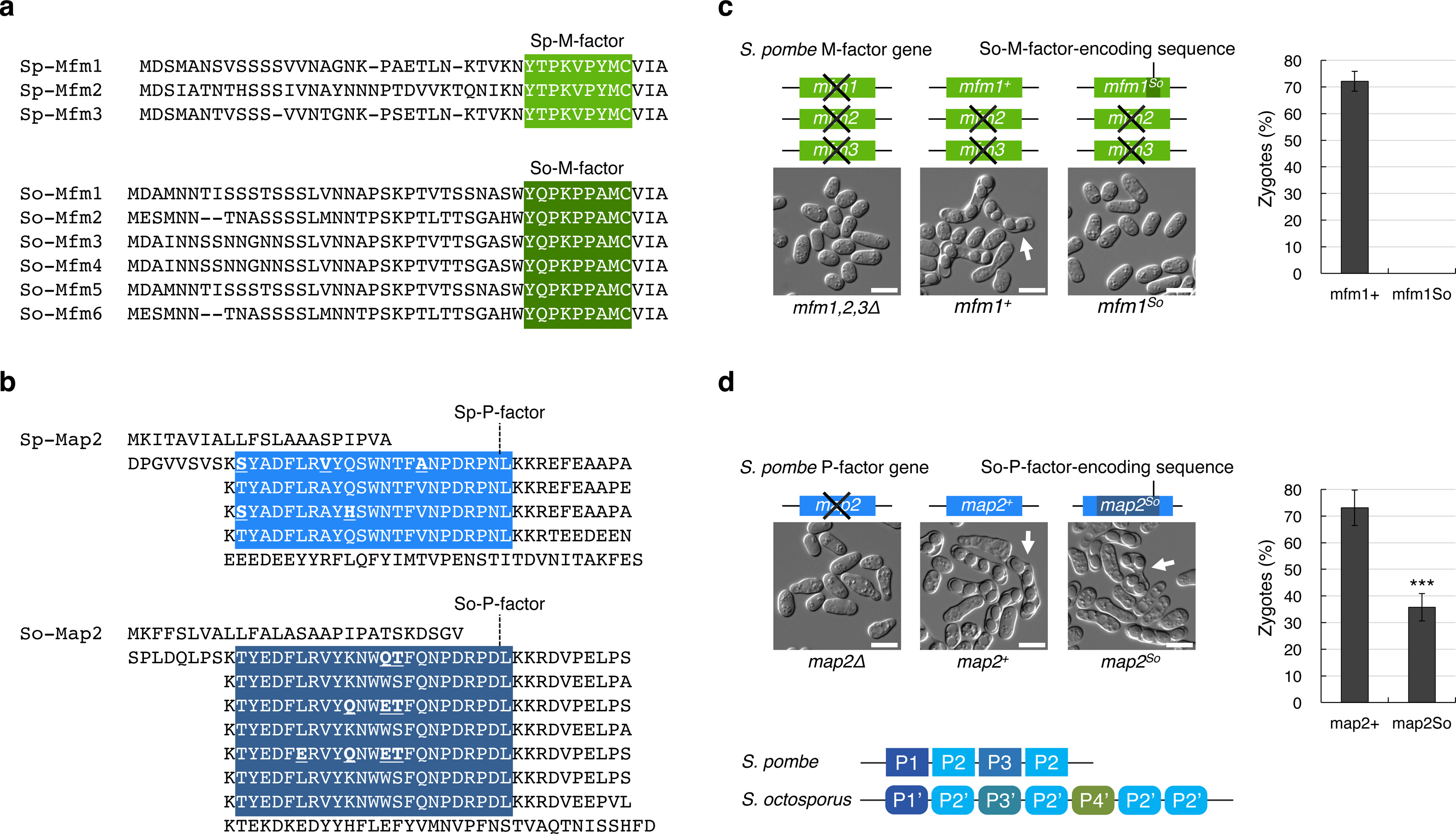
Comparison of mating pheromones between *S. pombe* and *S. octosporus*. **a**, Amino acid sequence alignment of precursors derived from the redundant genes encoding M-factor in *S. pombe* (3 genes) (top), and So-M-factor in *S. octosporus* (6 genes) (bottom). M-factor precursor proteins of *S. octosporus* are tentatively termed So-Mfm1-So-Mfm6. **b**, Amino acid sequence alignment of precursors derived from the gene encoding P-factor in *S. pombe* (top), and So-P-factor in *S. octosporus* (bottom). The P-factor precursor protein of *S. octosporus* is tentatively termed So-Map2. The native *So-map2* gene contains seven tandem So-P-factor-encoding repeats (P1’-P2’-P3’-P2’-P4’-P2’-P2’). Multiple sequence alignment was done by a standard algorithm. The mature peptide sequences are indicated by white letters on a color background. The P2’ peptide is the standard So-P-factor of *S. octosporus;* amino acids that differ from the standard peptides are underlined in bold. **c**, Mating efficiency of the *S. pombe* strain producing only So-M-factors. The strain in which the native *mfm1*^+^ gene was replaced with the *mfm1^So^* gene was sterile. **d**, Mating efficiency of the *S. pombe* strain producing only So-P-factors. The strain in which the native *map2*^+^ gene was replaced with the *map2^So^* gene showed partial fertility (zygotes (%), 35.8 ± 5.1%), as compared with the wild type (zygotes (%), 73.1 ± 6.7%). Data are the mean ± S.E.M. of triplicate samples. Statistical significance was assessed by *t*-test (\*\*\**p* <0.001). Typical images of mating cells (arrows, tetrad) are shown. Scale bar, 5 μm.

### P-factor is interchangeable between fission yeast species, but M-factor is not

Next, we assessed whether the pheromone peptides of *S. octosporus* are effective on *S. pombe* cells. First, the wild-type *S. pombe mfm1*^+^ gene was integrated into the genome of an M-factor-less strain (FY23412; Supplementary Table 1), which led to high mating efficiency (72.1 ± 3.7%) (Fig. 5c). Next, we replaced the Sp-M-factor-encoding sequence in the *mfm1*^+^ gene with the So-M-factor-encoding sequence (*mfm1^So^*). The resulting strain, which produced only So-M-factor instead of Sp-M-factor, was found to be completely sterile (Fig. 5c). In short, *S. pombe* P-cells were unable to mate with M-cells producing So-M-factor. By contrast, replacement of the Sp-P-factor-encoding sequence (4 repeats) in the *map2*^+^ gene with the So-P-factor-encoding sequence (7 repeats) (*map2^So^*), resulted in a strain with approximately half the mating frequency of the strain with the *map2*^+^ gene (35.8 ± 5.1%) (Fig. 5d). Thus, *S. pombe* M-cells can mate with P-cells that produce So-P-factors, indicating that at least one of the So-P-factors is effective on *S. pombe* cells. We also replaced the So-P-factor-encoding sequence (7 repeats) in the *So-map2*^+^ gene with the Sp-P-factor-encoding sequence (4 repeats) (*So-map2^Sp^*) in *S. octosporus* cells. The resulting strain, which produced only Sp-P-factor instead of So-P-factor, retained mating ability (17.3 ± 4.9%; see Supplementary Fig. 3). Thus, at least one of the Sp-P-factors is also effective on *S. octosporus* cells.

Three of the nine amino acids of So-M-factor were found to differ from those of Sp-M-factor (Fig. 6a). To assess whether So-M-factor is recognized at all by the M-factor receptor of *S. pombe*, we carried out a shmooing assay (see Fig. 4) in which we measured the L/W ratio of cells treated with synthetic So-M-factors at various concentrations in nitrogen-free liquid medium (EMM2-N). The M-factor-sensitive strain (TS405; Supplementary Table 1) significantly induced cell elongation after 1 day of incubation with 100 nM Sp-M-factor, but showed no elongation when treated with So-M-factor (Fig. 6b). Under these conditions, approximately 40% of cells treated with Sp-M-factor peptide underwent shmoo formation, whereas no cells treated with So-M-factor peptide underwent shmooing (Fig. 6b). When the concentration of So-M-factor was raised to 1000 nM, the cells underwent shmoo elongation a little (Fig. 6b). These results imply that *S. pombe* is reproductively isolated from *S. octosporus* owing to the lack of compatibility of M-factor.

**Figure 6.**
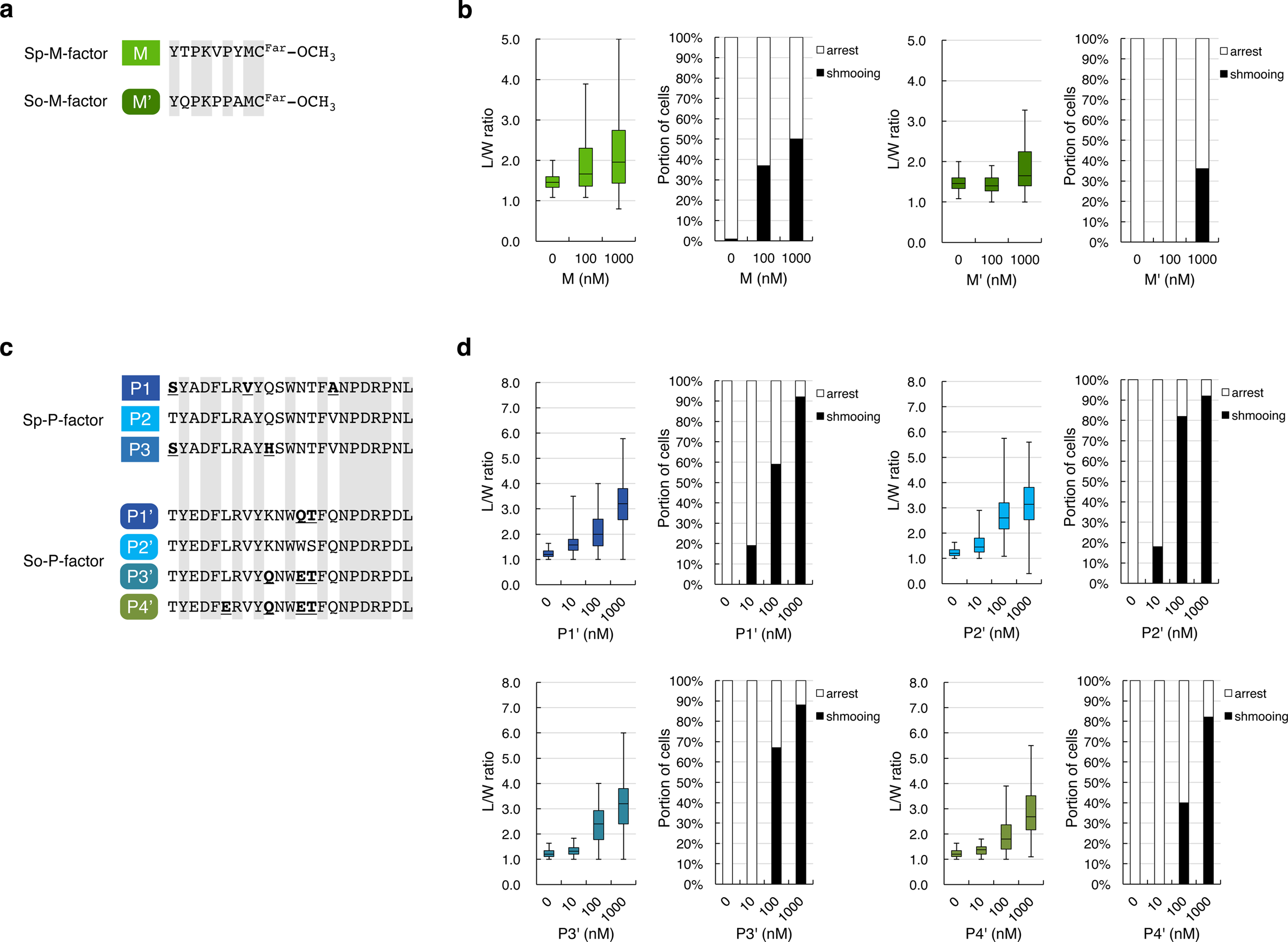
Effects of mating pheromones of *S. octosporus* on *S. pombe* cells. **a**, Comparison of the amino sequences of Sp-M-factor and So-M-factor. Identical amino acids are shown in gray, indicating that three of the nine amino acid residues (T2, V5, and Y7) of M-factor differ between the two species. **b**, Shmooing assay of synthetic So-M-factor peptide and *S. pombe* cells. P-cells lacking the *rgs1*^+^ (TS405) were treated with synthetic M-factor at different concentrations (0, 100, and 1000 nM) and incubated in EMM2-N medium with gentle shaking for 24 hours. The ability of each M-factor peptide to induce shmooing was assessed by the L/W ratio of a cell, as described in Fig. 4. **c**, Comparison of the amino sequences of Sp-P-factors and So-P-factors. Identical amino acids in all peptides are shown in gray, indicating that about 8 of the 23 amino acid residues of P-factor differ between the two species. The amino acids that differ within each species are underlined in bold. **d**, Shmooing assay of synthetic So-P-factor peptide and *S. pombe* cells. M-cells lacking the *sxa2*^+^ (TS402) were treated with synthetic So-P-factor at different concentrations (0, 10, 100, and 1000 nM) and incubated in EMM2-N medium with gentle shaking for 24 hours. The shmooing assay was evaluated as described in Fig. 4.

On average, 8 of the 23 amino acids of P-factor differed between the two species. To test whether the *S. octosporus* P-factor peptides (hereafter named P1’–P4’; Fig. 6c) are recognized by Mam2 of *S. pombe* (Sp-Mam2), we treated TS402 with each of the four synthetic So-P-factor peptides. Remarkably, cells formed visible shmoos in all cases (Fig. 6d). The most effective peptide was P2’; that is, one-fifth of *S. pombe* cells treated with P2’ peptide (10 nM) elongated a shmoo in response to P-factor after 1 day, and the proportion of shmooing cells increased to about 80% at a peptide concentration of 100 nM (Fig. 6d). In contrast, recognition of the P4’ peptide by Sp-Mam2 was relatively low. Taken altogether, these findings indicate that all of the So-P-factors have a partial effect on *S. pombe* M-cells.

### A pheromone-degrading enzyme might facilitate pheromone diversity

The above findings showed that *S. pombe* cells can respond to So-P-factors produced by *S. octosporus*. We therefore examined whether Sxa2 of *S. pombe* can degrade So-P-factors. Each of the synthetic So-P-factor peptides was mixed at 200 μM with culture supernatant containing active Sxa2, and leucine concentration was measured at fixed time points, as described above for Sp-P-factors. All four So-P-factor peptides (P1’-P4’) were considerably degraded in the presence of Sxa2 (Table 2). The P2’ peptide was degraded most efficiently, with 83.7 ± 4.0 μM degraded after 60 minutes of incubation; as compared with Sp-P-factor peptides, however, Sxa2 inefficiently removed the Leu residue from So-P-factor peptides (Table 2). In conclusion, Sxa2 acts most effectively on Sp-P-factors but has the capacity to degrade P-factor peptides from related species such as *S. octosporus*, probably limiting the interspecies mating response.

Lastly, we examined the diversity of Sxa2 in nature. In the 150 *S. pombe* wild strains, there were four different amino acid sequences of Sxa2 (Table 1 and Supplementary Table 4). In exploring the asymmetric diversification of the two mating pheromones, it is notable that M-factor is not degraded by a specific enzyme. We speculate that overlapping regions of P-factor (probably the C terminus) are recognized by both its receptor Mam2 and its degradation enzyme Sxa2, thereby facilitating coevolution of their substrate specificities together with asymmetric divergence of P-factor (Fig. 7).

**Figure 7.**
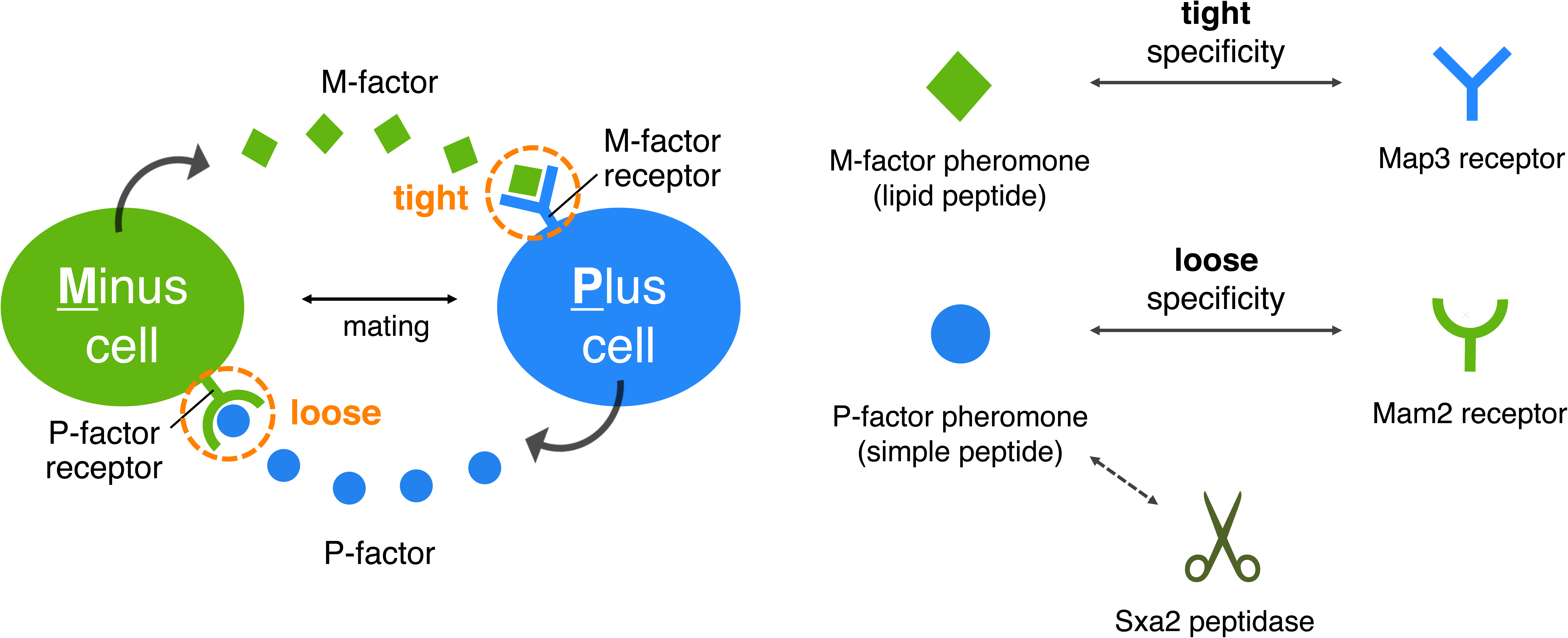
An asymmetric system of pheromone recognition in fission yeast. The specificity of recognition of M-factor (lipid peptide) is extremely tight, whereas that of the P-factor (simple peptide) is relatively loose, allowing cross-reactions to occur between two yeast species. This possession of two pathways for the mating pheromones might guarantee not only tight recognition of mating partners, but also flexible adaptation to various mutational changes in a combination of a pheromone/receptor. M-factor interacts with only Map3, but P-factor interacts with both Sxa2 and Mam2. Therefore, the diversity of the two mating pheromones might facilitate asymmetrically in nature.

## Discussion

Mating in yeasts critically depends on molecular recognition of two peptidyl mating pheromones by their corresponding receptor on each mating partner. Therefore, mutational alterations of the pheromone–receptor system affect the recognition between partners, resulting in prezygotic isolation. In this study, we explored the coevolution of the pheromones and cognate receptors of *S. pombe* in nature. We found that the amino acid sequences of M-factor and its receptor Map3 are completely conserved, whereas those of P-factor and its receptor Mam2 are very diversified (Table 1). Such asymmetric diversification of the two mating pheromones was also seen in the related species *S. octosporus* (Fig. 5a, b). Moreover, we noticed that So-M-factor was not functional on *S. pombe* cells, whereas all of the So-P-factors tested were partially functional; in other words, *S. pombe* cells were capable of mating with cells producing only So-P-factors (Fig. 5c, d). Therefore, it is more likely that the recognition specificities of the two pheromone receptors vary in strictness for their respective pheromones. Map3 and Mam2 are both class IV GPCRs, but differ clearly in their primary structure. Map3 has high homology to Ste3 (a GPCR for a-factor in *S. cerevisiae*), whereas Mam2 is similar to Ste2 (a GPCR for α-factor in *S. cerevisiae^39^*), indicating that two different types of pheromone receptor are commonly conserved in fungi. Recently, Rogers *et al.^16^* reported that a-type pheromones (lipid peptide) either promote efficient mating completely or do not promote it not at all, while α-type pheromones (simple peptide) show a more graded distribution of mating efficiency in *S. cerevisiae*. Thus, the different specificities of the two GPCRs might lead to asymmetric competence and diversification of pheromones.

*S. pombe* has three genes encoding Sp-M-factor, and *S. octosporus* has six putative genes encoding So-M-factor (Fig. 5a). Such redundancy might enable the cells to alter one copy of M-factor to adapt to genetic changes in Map3, while keeping the others unchanged. In fact, the N-terminal half of M-factor has been shown to be dispensable for recognition by Map3^36^. Nevertheless, the redundant genes encoding M-factor peptides generate the same sequence of amino acids in nature (Fig. 5a). We considered that there are two potential explanations for this: first, the amount of mature M-factor might affect mating frequency; second, mutation of M-factor might have a dominant-negative effect. To examine the first possibility, we constructed strains harboring different number of *mfm* genes (TS250–TS255; Supplementary Table 1) and then examined the frequency of zygotes. As the number of *mfm* genes decreased, the mating frequency also decreased, but only a little (Supplementary Fig. 2a). This is consistent with previous data (despite different experimental conditions), of Nielsen *et al.^40^*, who reported that any one of the three genes of M-factor is sufficient for mating, and all genes are able to produce active M-factor peptide in the wild-type strain. However, this might not reflect the actual situation in nature. For example, efficient copulation might depend largely on the number of M-factor genes under more severe nutrient-limited conditions. We could not confirm whether there is an increase or decrease in *mfm* genes in the 150 wild *S. pombe* strains in this study, but the importance of *mfm* gene number will be a topic for future study. To examine whether mutated M-factor has a dominant-negative effect, we inspected the frequency of zygotes of some strains producing two different M-factor peptides (TS340–TS343; Supplementary Table 1). Production of any multiple M-factors did not have a marked effect on mating efficiency (Supplementary Fig. 2b) as far as we investigated. These results indicate that mutated M-factor peptide is unlikely to prevent the activity of the wild-type itself. At present, therefore, it is unclear why the amino acid sequences of M-factor and Map3 are strictly conserved in nature.

In the *S. pombe* wild strains, the number of P-factor-encoding repeats in the *map2* gene varied from 4 to 8 (Fig. 2a). Variations in repeat number in the *Saccharomyces* genus have previously been reported^41-44^. Decreasing numbers of repeats in Mfαl, a structural gene for α-factor pheromone, results in a stepwise decrease in α-factor production in *Saccharomyces cerevisiae^45^*. Rogers *et al*.^44^ also showed that experimentally varying the repeat number in the Mfαl gene affects the amount of a-factor in *Saccharomyces paradoxus*. Our experimental data revealed that removing a single repeat of the 4-repeat sequence in *map2* has a marked effect on mating frequency (Fig. 2b). A high level of P-factor is likely to be required to overcome Sxa2 activity. In addition, a previous study suggested that the secretion pathway has a minimum size requirement for transportation^46^. Hence, the product of the *map2* gene might need to have sufficient length for processing by the biosynthetic pathway. Increasing the number of repeats enables the cells to avoid copulation failure, but may not provide a benefit for mating (Supplementary Fig. 1a). For example, Rogers *et al.^44^* also showed that an 8-repeat strain is more unfavorable than a 6-repeat strain for mating choice in *S. cerevisiae*, probably due to reduced pheromone production caused by a decrease in the rate of translation. When the various *map2* genes (4–8 repeats) obtained from the 150 wild strains were integrated into the FY23418 strain, almost all of the resulting strains showed high mating frequency (Supplementary Fig. 1a). The only exception was the *map2* gene (6 repeats) obtained from the 22CBS10468 strain (*map2_D8;* Supplementary Table 3), where a spacer sequence located between the fourth and the fifth repeat was changed from KKR to KKC (Supplementary Table 4). The kexin-related endopeptidase Krp1 cleaves the KKR motif (three basic amino acids) in the Golgi during the biosynthetic pathway to generate mature P-factor^47^; therefore, such a mutation might affect the production of P-factor. A relationship between repeat number and sexual fitness in yeast has been also reported. Yeast cells choose a favorable partner producing the highest levels of pheromone, whereas a cell that cannot produce pheromones is not chosen as a mating partner by the opposite mating cell^48-50^. Such fluctuations in the repeat number of P-factor are likely to have an influence on various factors related to mating events.

All of the wild *S. pombe* strains produced at least three different P-factors as far as we investigated (Fig. 2a). Curiously, however, our experimental data clearly showed that the P2 peptide was much more efficiently recognized by Mam2 as compared with the others *in vivo* and *in vitro* (Figs. 2c and 4). Why don’t *S. pombe* cells produce only P2 peptides? Interestingly, we found that cells producing multiple different P-factors seemed to be slightly preferred as a mate by the opposite cell type (Fig. 3). This might be explained by the differences in the extent of degradation by Sxa2. That is, although P2 peptide is recognized more efficiently by native Mam2, it is also degraded by Sxa2 more rapidly (Table 2). We speculate that the production of different peptides, which are more resistant to degradation, might be helpful to avoid the risk of their complete breakdown. We further noticed that there is a positive relationship between the recognition of P-factor by Mam2 and its efficient degradation by Sxa2 (Fig. 4 and Table 2). This is because, both Mam2 and Sxa2 are likely to depend on the same regions of P-factor activity. A recent study also suggests that coevolution of Ste2 (a receptor for α-pheromone) and Bar1 (a peptidase of α-pheromone) can occur, together with evolution of α-pheromone in *Candida Albicans*, because Ste2 and Bar1 recognize the overlapping regions of α-pheromone^51^. In this study, different polymorphisms of Mam2 and Sxa2 were identified in wild *S. pombe* strains (Table 1). Although we did not obtain conclusive evidence that novel compatible combinations of the P-factor/Mam2 and P-factor/Sxa2 pairs can occur (Supplementary Fig. 1c, d), such coevolution might proceed little by little, even though reproductive isolation would not be prevented during this time.

In *S. pombe*, M-factor is a farnesylated hydrophobic peptide, whereas P-factor is an unmodified hydrophilic peptide. This asymmetry in the chemical properties of the two pheromones is conserved across ascomycetes including yeasts^13^. Notably, in *S. cerevisiae*, it is known that a simple peptide α-factor promotes secretion of a lipid peptide a-factor, but a-factor is less important for secretion of α-factor^52^. It might have significant benefits for mating process in yeasts that production of the pheromone with high specificity (lipid peptide) is promoted by the pheromone with relatively low specificity (simple peptide). A hydrophilic peptide is more diffusible, because it might reach far-away cells, likely enabling to become aware of the existence of mating partners rapidly. Our findings lead us to hypothesize that 1) the sexual behavior of individual species in nature is controlled by sexual interactions across species, 2) the farnesyl group of pheromone is the key determinant for mating partner discrimination. Probably, the asymmetric system that has evolved in yeast for recognition of the two pheromones might allow flexible adaptation of the simple peptide to mutational changes in the pheromone/receptor pair, while maintaining tight recognition for mating partners by the lipid peptide (Fig. 7), which is likely to be the main mechanism underlying the process of reproductive isolation leading to speciation in yeasts.

## Methods

### Strains, media, and culture conditions

The strains constructed in this study are listed in Supplementary Table 1. The 150 wild *S. pombe* strains with a different origin from the standard laboratory strain (Leupold’s strain, L968^33^) were obtained from the National BioResource Project (deposited by J. Kohli, M. Sipiczki, G. Smith, H. Levin, A. Klar, and N. Rhind), H. Innan^34^, and J. Bähler^35^ (Supplementary Table 2).

Cells were grown in yeast-extract (YE) medium (0.5% Bacto yeast extract and 3.0% D-glucose) supplemented with adenine (75 mg/l), uracil (50 mg/l) and leucine (50 mg/l). For solid medium, 1.5% agar was added to YE medium (YEA). Where appropriate, antibiotics (G418, Hygromycin B, and nourseothricin) were added to YEA at a final concentration of 100 μg/ml. Edinburgh minimal medium 2 (EMM2)^53^ was also used for growth. Synthetic Dextrose (SD) medium (0.67% Bacto yeast nitrogen base without amino acids, 2.0% D-glucose, and 1.5% agar) was used to select auxotrophic mutants of *S. pombe*. Malt-extract agar (MEA) medium (3.0% Bacto Malt Extract and 1.5% agar, pH 5.5), Edinburgh Minimal Medium without NH_4_Cl (EMM2-N) medium^53^, and Pombe Minimal Glutamate (PMG) medium^54^ were used for mating and sporulation. Cells were grown and conjugated for a few days at 30°C.

### Sequencing analysis of wild strains

Genomic DNA was extracted from overnight cultures grown in YE medium. All PCR reactions were carried out using KOD-FX Neo (TOYOBO). Each of the DNA fragments containing *mfm1, mfm2, mfm3, map2, map3, mam2*, and *sxa2* was amplified by the following primer sets: oTS83/84, oTS507/508, oTS509/510, oTS81/82, oTS91/92, oTS89/90, and oTS624/625, respectively (all primers are listed in Supplementary Table 5). The resulting DNA fragments were purified through a Wizard^®^ PCR cleaning system (Promega). The PCR products were subjected to nucleotide sequencing by using a 3130xl Genetic Analyzer (Applied Biosystems) and the internally specific primers: oTS504 (*mfm1*), oTS505 (*mfm2*), oTS506 (*mfm3*), oTS85/86 (*map2*), oTS157/158 (*map3*), oTS151/152 (*mam2*), and oTS647/648/658 (*sxa2*). The sequences obtained were compared with the corresponding sequence of L968. Differences from the L968 sequence are listed in Supplementary Tables 3 and 4.

### Construction of strains with variable numbers of P2 peptide in the Map2 ORF

The *map2*^+^ gene (~2.6 kb) containing its promoter and terminator regions was amplified from L968 genomic DNA by the primer set oTS81/82. The DNA fragment was fused by using a Gibson Assembly (New England Biolabs) to a linearized vector derived from the integration vector pBS-ade6^36^ which was prepared by inverse PCR with the primer set oTS79/80. The resultant plasmid, pTS13 (all plasmids used in this study are listed in Supplementary Table 6), was used as a starting template. To create a P2 unit, the correct two oligos (oTS73 and oTS93; reverse complements of each other) were mixed in Annealing Buffer (10 mM Tris-HCl, pH 7.4, 1 mM EDTA, and 50 mM NaCl), incubated at 70°C for 10 minutes by a GeneAmp® PCR System 9700, and then slowly cooled to 25°C (~1 hours). The resulting double-strand DNA fragment (the P2 unit) was purified as described above. The DNA fragment that was amplified from the P2 unit by the primer set oTS74/75 was fused to a linearized vector derived from pTS13, which was prepared by inverse PCR with the primer set oTS71/72 to replace the *map2*^+^ gene. The resultant plasmid was referred to as pTS10.

To create Map2 ORFs with 2–4 P2 repeats, a further three P2 units carrying slightly different sequences at both sides were amplified from pTS10 by the appropriate primers (see Supplementary Table 7 for all combinations of primers used for P-factor-related plasmids). Three mixed DNA fragments were fused to a linearized vector derived from pTS10, which was prepared by inverse PCR with the primer set oTS72/179. The resultant plasmids had various random repeats in the Map2 ORF. The number of repeats in each plasmid was checked by PCR using oTS85/86, and plasmids with the desired number of repeats were sequenced to confirm their sequences. Thus, pTS47, pTS48, and pTS66 were constructed. The plasmids were cut near the center of the *ade6+* gene with *Bam*HI, and integrated at the *ade6* locus on chromosome III in the FY23418 strain.

### Construction of strains producing various P-factor peptides (4 peptide repeats)

To generate pTS54, pTS55, pTS145, pTS146, and pTS233, inverse PCR was carried out using pTS10 and the primer sets oTS202/203, oTS204/205, oTS401/402, oTS403/404, and oTS204/404, respectively. The amplified PCR products were subjected to *Dpn*I treatment, and 5′-phosphorylated by T4 polynucleotide kinase (TaKaRa), ligated by T4 ligase (TaKaRa), and transformed into *Escherichia coli* (DH5α). The introduced mutations were confirmed by sequencing the recovered plasmids.

All plasmids carrying 4-repeat Map2 ORFs were constructed in the following way. Three DNA fragments encoding P-factor units carrying slightly different sequences at both sides were amplified from a template plasmid by the appropriate primers (see Supplementary Table 7). Each of the three mixed DNA fragments was fused to the corresponding linearized vector, which was prepared by inverse PCR with the appropriate primers. The number of repeats in the resultant plasmids was checked by PCR using the primer set oTS85/86, and plasmids with 4 repeats were sequenced to confirm their sequences. In addition, the *map2* gene (~2.6 kb) containing the promoter and terminator regions was amplified from the genomic DNA of nine wild strains (01CBS10391, 04CBS2775, 06CBS10460, 13CBS10504, 22CBS10468, 24CBS5682, 25CBS5680, 32CBS352, and 55FY28965) by the primer set oTS81/82. Each DNA fragment was fused to a linearized vector derived from the integration vector pBS-ade6, which was prepared by inverse PCR with the primer set oTS79/80. All of the obtained plasmids were integrated into the FY23418 strain as described above.

### Quantitative assay of zygote formation

Cells grown on YEA overnight were resuspended in sterilized water to a cell density of 1 × 10^8^ cells/ml. A 30-μl aliquot of the resulting suspension was spotted onto sporulation media (MEA for *S. pombe;* PMG for *S. octosporus*), and then incubated for 2 days (S. *pombe*) or 3 days (S. *octosporus*) at 30°C unless stated otherwise. Cells were counted under a differential interference contrast (DIC) microscope. Cell types were classified into four groups: vegetative cells (V), zygotes (Z), asci (A), and free spores (S). The percentage of zygotes was calculated by using the following equation^36,55^.

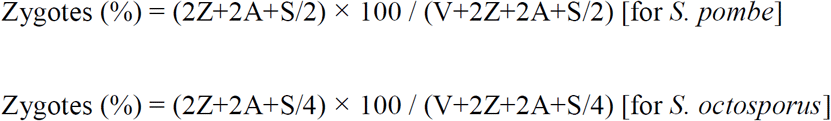

In all cases, triplicate samples (at least 300 cells each) were counted, and the mean and standard error of mean (S.E.M.) were calculated.

### Competitive test of hybrid formation by recombinant frequency

Heterothallic haploid strains each carrying a chromosomal drug-resistance marker (kanMX6, hphMX6, or natMX6) were cultured on YEA overnight, and the same cell numbers of the M-strain and two competing P-strains were mixed in sterilized water. A 30-μl aliquot of the resulting suspension was spotted onto MEA, and incubated for exactly 24 hours at 30°C. The mixed cells were allowed to mate, and the resulting hybrid diploids were sporulated to form spores. The cell suspension was diluted and spread on YEA plates containing different combinations of drugs. The number of colonies was counted after 3 days of incubation at 30°C. The ratio of recombinant frequency was calculated by determining the number of doubly-resistant colonies, which was normalized by the colony numbers on YEA plates containing either one of the two drugs. Three separate competitive tests were carried out.

### Shmooing assay

P-factor and M-factor peptides were chemically synthesized (Eurofins) for the shmooing assay. The purity of the preparations was over 95% (HPLC). P-factor was dissolved in dimethyl sulfoxide (DMSO) at a concentration of 500 μM, and M-factor was dissolved in methanol (MeOH) at a concentration of 1 mM. The stock solutions were diluted with culture medium to the appropriate dilution ratio. For the assay, heterothallic haploid cells were grown in YE medium overnight, washed with sterilized water three times, and then resuspended in EMM2-N medium at a cell density of 4 × 10^7^ cells/ml. The cells were treated with the synthetic pheromone, and incubated for exactly 24 hours with gentle shaking.

To assess whether Mam2 could recognize different P-factors, a P-factor sensitive strain lacking the *sxa2*^+^ gene (TS402) was used. The cells were incubated with synthetic P-factors at different concentrations (0, 10, 100, and 1000 nM) for 24 hours and observed by a DIC microscope. Images were recorded, and the length (L) and width (W) of a cell were measured to determine the L/W ratio. Cells with an L/W ratio of 2.0 or more were defined as shmooing cells; other cells were defined as arrested. In all cases, at least 100 cells each were measured. In the reciprocal experiment, an M-factor-sensitive strain lacking the *rgs1*^+^ gene^56^ (TS405) was used to assess whether Map3 can recognize different M-factors. The cells were treated with synthetic M-factors at different concentrations (0, 100, and 1000 nM) for 24 hours, and analyzed as described above.

### Preparation of culture medium from cells expressing Sxa2

The pREP vector was used to express *sxa2*^+^ under the control of the thiamine-repressible *nmt1*^+^ promoter in *S. pombe^37^*. First, a KanMX6 cassette amplified from pFA6a-kanMX6^57^ by the primer set oTS299/300 was fused to the linearized pREP1 vector prepared by inverse PCR with the primer set oTS301/302. In the resultant plasmid pTS111, the *LEU2* gene was replaced with a KanMX6 cassette. Next, the *sxa2*^+^ gene (~1.5 kb) was amplified from L968 genomic DNA by the primer set oTS729/730, and fused to a linearized vector derived from the integration plasmid pTS111, which was prepared by inverse PCR with the primer set oTS305/306. The resultant plasmid, pTS284, was transformed into TS402. Thus, a heterothallic strain (TS407) ectopically overexpressing *sxa2*^+^ under the control of the *nmt1*^+^ promoter was obtained.

The TS407 strain was precultured overnight in YE medium containing G418, and then washed with sterilized water three times. Cultures were inoculated into EMM2 medium containing G418 without thiamine (to induce the *nmt1*^+^ promoter) at a cell density of 1 × 10^7^ cells/ml, and incubated with aeration for 2 days. The cell culture supernatant was passed through a 0.22-μm filter (Merck Millipore) to completely remove cell debris, and then concentrated 20-fold by ultrafiltration through a Vivaspin® 20-10K (GE healthcare), because ultrafiltration has been shown to concentrate the carboxypeptidase without loss of Sxa2 activity^29^. Thus, culturemedium including active Sxa2 was obtained. As a control, pTS111 (*sxa2^−^*) was transformed into TS402, and the same preparation of culture medium was carried out.

### *In vitro* assay of the degradation level of P-factor

All reactions were performed at 30°C in 50 mM citrate buffer, pH 5.5, with 200 μM synthetic P-factor as described previously^58^. Culture medium containing 100 ng of protein, as determined by a Bradford protein assay (BioRad), from strain TS406 or TS407 was added to a solution of P-factor, and then incubated for appropriate times (0, 10, and 60 mins) with gentle shaking via the constant-temperature incubator shaker MBR-022UP (TAITEC). Reactions were stopped by adding trifluoroacetic acid to 0.5%. Leucine concentration in the samples was measured by using a Branched Chain Amino Acid (BCAA) Assay kit (Cosmo Bio co.) in accordance with the manufacturer’s protocol and a Model 680 Microplate Reader (BioRad). Carboxypeptidase assays were performed in at least triplicates, and the mean ± S.E.M. was calculated. As a negative control, it was verified that almost no leucine was detected in an assay with a P-factor peptide lacking the C-terminal (Table 2).

### Construction of strains producing pheromones from different species

Site-directed mutagenesis of the M-factor-coding gene, *mfm1*^+^, was conducted by using the *in vitro* mutagenesis reagents Quik®-Change (Stratagene) as previously reported^36^. In briefly, DNA replication using pBS-ade6(*mfm1*^+^)^36^ as a template to generate a mutated strand was performed in accordance with the manufacturer’s protocol. Primers were designed to introduce the desired mutated triplet codon at a specific site of the *mfm1*^+^ ORF. After replication, the reaction mixture was treated with *Dpn*I to digest the *mfm1*^+^ strand. The *Dpn*I-treated plasmid was then transformed into DH5α. The introduced mutation was confirmed by sequencing the recovered plasmids. Next, the resultant plasmids pTS189 was integrated at the *ade6* locus of an M-factor-less strain (FY23412). The strain expressed only the mutated *mfm1* gene and thus produced So-M-factor. By contrast, the *So-map2*^+^ gene lacking its signal sequence and two *N*-linked glycosylation sites was amplified from yFS286 genomic DNA by the primer set oTS57/58. The DNA fragment was fused to a linearized vector derived from pTS13, which was prepared by inverse PCR with the primer set oTS55/56 to replace the *map2*^+^ gene. The resultant plasmid, pTS15, was integrated into the FY23418 strain as described above. This strain produced only So-P-factors, and no Sp-P-factors.

Next, we constructed an *S. octosporus* strain producing only Sp-P-factors. To delete the native *So-map2*^+^ gene, the 3’-downstream sequence (1 kb) was first amplified from yFS286 genomic DNA by using the primer set oTS47/48. The DNA fragment was fused to a linearized vector derived from pFA6a-kanMX6, which was prepared by inverse PCR with the primer set oTS37/38 to construct the plasmid pTS5. Next, the 5’-downstream sequence (1 kb) was amplified from yFS286 genomic DNA by using the primer set oTS46/78. The DNA fragment was fused to a linearized vector derived from pTS5, which was prepared by inverse PCR with the primer set oTS35/36 as described above. Lastly, pTS11 was constructed. The DNA fragment from this plasmid, namely, a 3.5-kb fragment amplified from pTS11 by using the primer set oTS48/78, was purified, and 1 μg of the DNA was transformed into yFS286 by our previously described method^55^. The *So-map2*^+^ gene was successfully disrupted by a homologous recombination, and the resultant strain TS12 was sterile.

The *So-map2*^+^ gene (~3.3 kb) containing the promoter and terminator regions was amplified from yFS286 genomic DNA by the primer set oTS78/626. The DNA fragment was fused to a linearized vector derived from the integration vector pFA6a-natMX6, which was prepared by inverse PCR with the primer set oTS35/36 to construct the plasmid pTS247. The *Sp-map2*^+^ gene lacking its signal sequence and two *N*-linked glycosylation sites was amplified from L968 genomic DNA by the primer set oTS635/636. The DNA fragment was fused to a linearized vector derived from pTS247, which was prepared by inverse PCR with the primer set oTS633/634 to replace the *So-map2*^+^ gene. The resultant plasmid pTS250 was integrated into the TS12 strain after restriction with *Bam*HI near the center of the terminator region. This strain produced only Sp-P-factors, and no So-P-factors.

### Construction of strains expressing various *mam2* genes

The *mam2*^+^ gene (~3.1 kb) containing its promoter and terminator regions was amplified from L968 genomic DNA by the primer set oTS90/99. The DNA fragment was fused to a linearized vector derived from the integration vector pFA6a-hphMX6, which was prepared by inverse PCR with the primer set oTS35/36 to construct the plasmid pTS14. The *mam2* gene (~1.0 kb) was amplified from the genomic DNAs of four wild strains (02CBS2628, 04CBS2775, 06CBS10460, and 26CBS5557) by the primer set oTS586/587. Each of the DNA fragments was fused to a linearized vector derived from pTS14, which was prepared by inverse PCR with the primer set oTS149/150 to replace the *mam2*^+^ gene. All obtained plasmids were integrated into the FY12677 strain after restriction with *Afe*I near the center of the terminator region.

### Construction of strains expressing various *sxa2* genes

The *sxa2*^+^ gene (~3.5 kb) containing its promoter and terminator regions was amplified from L968 genomic DNA by the primer set oTS624/625. The DNA fragment was fused to a linearized vector derived from the integration vector pFA6a-natMX6, which was prepared by inverse PCR with the primer set oTS35/36 to construct the plasmid pTS246. The *sxa2* gene (~1.5 kb) was amplified from the genomic DNAs of three wild strains (01CBS10391, 04CBS2775, and 101FY29038) by the primer set oTS664/665. Each of the DNA fragments was fused to a linearized vector derived from pTS246, which was prepared by inverse PCR with the primer set oTS627/628 to replace the *sxa2*^+^ gene. All obtained plasmids were integrated into the TS356 strain after restriction with *Bam*HI near the center of the terminator region.

## Acknowledgments

We thank J. Kohli, M. Sipiczki, G. Smith, H. Levin, A. Klar (deceased), N. Rhind, H. Innan, T. Iida, the National BioResource Project (NBRP) and the Japan Collection of Microorganisms (JCM) for strains, and Y. Kurita at National Institute of Genetics for sequence analyses. This work was supported by Grant-in-Aid for JSPS Research Fellow (Grant Number JP15J03416) to T.S., Grant-in-Aid for Young Scientists (B) (Grant Number JP17K15181) to T.S., the Sumitomo Foundation (No. 160924) to T.S., and Dr. Yoshifumi Jigami Memorial Fund, The Society of Yeast Scientists to T.S.

## Author contributions

T.S. designed and performed most experiments. C.S. helped for sequence analyses and provided discussion in the early stages of this work. H.N. supervised this project. T.S. drafted the manuscript with contributions from C.S. and H.N.

## Supplementary information

**Figure S1 Mating efficiency of various strains. a**, Zygote frequency of strains with a *map2* ORF containing different numbers of P-factor-encoding repeats: WT (4 repeats, L968), D9 (4 repeats, 24CBS5682), D13 (4 repeats, 55FY28965), D11 (5 repeats, 32CBS352), D1 (5 repeats, 01CBS10391), D5 (5 repeats, 06CBS10460), D8 (6 repeats, 22CBS10468), D3 (6 repeats, 04CBS2775), D10 (7 repeats, 25CBS5680), and D7 (8 repeats, 13CBS10504). **b**, Zygote frequency of strains producing various P-factors (4 repeats). **c**, Zygote frequency of strains expressing various *mam2* genes: WT (L968), F2 (04CBS2775), F1 (02CBS2628), F5 (26CBS5557), and F3 (06CBS10460). **d**, Zygote frequency of strains expressing various *sxa2* genes or lacking the *sxa2*^+^ gene: WT (L968), G1 (01CBS10391), G4 (04CBS2775), and G7 (101FY29038). At least 300 cells were examined for each sample. Data are the mean ± S.E.M. of triplicate samples. Statistical significance was assessed by *t*-test (\**p* <0.05, \*\*\**p* <0.001).

**Figure S2 Mating efficiency of strains carrying different number of *mfm* genes. a**, Zygote frequency of strains defective in *mfm1, mfm2*, and *mfm3*. **b**, Zygote frequency of strains producing two different M-factor peptides. A previous study reported that, in M-factor, T2Q substitution has no inhibitory effect, V5P substitution leads to severe inhibition, and Y7A substitution causes sterility. M-factor-T2Q,V5P,Y7A is equivalent to So-M-factor. Cells were incubated on MEA for 1 day at 30°C. Data are the mean ± S.E.M. of triplicate samples. Statistical significance was assessed by *t*-test (\**p* <0.05, \*\**p* <0.01, and \*\*\**p* <0.001).

**Figure S3 Effects of P-factors on *S. octosporus* cells.** Mating efficiency of an *S. octosporus* strain producing only P-factors was determined by the frequency of zygotes. The strain in which the native *So-map2*^+^ gene was replaced with the *So-map2^Sp^* gene showed partially fertility (zygotes (%), 17.3 ± 4.9%), but fertility was less than the strain with the wild-type gene. Data are the mean ± S.E.M. of triplicate samples. Statistical significance was assessed by *t*-test (\*\*\**p* <0.001). Typical images of mating cells (arrows, octad) are shown. Scale bar, 10 μm.

Table S1 Strains used in this study.

Table S2 List of *S. pombe* wild strains whose origin differs from L968.

Table S3 Nucleotide sequence pattern of seven pheromone-associated genes in 150 *S. pombe* wild strains.

Table S4 Polymorphic alleles in the seven pheromone-associated genes of *S. pombe* wild strains.

Table S5 Primers used in this study.

Table S6 Plasmids used in this study.

Table S7

